# TMEM55A-mediated PI5P signaling regulates α-cell actin depolymerization and glucagon secretion

**DOI:** 10.1101/2024.12.16.628242

**Authors:** Xiong Liu, Theodore dos Santos, Aliya F. Spigelman, Shawn Duckett, Nancy Smith, Kunimasa Suzuki, Patrick E. MacDonald

**Affiliations:** Department of Pharmacology, University of Alberta, Edmonton, AB T6G 2E1, Canada; Alberta Diabetes Institute, University of Alberta, Edmonton, T6G 2E1, Canada

## Abstract

Diabetes is associated with the dysfunction of glucagon-producing pancreatic islet α-cells, although the underlying mechanisms regulating glucagon secretion and α-cell dysfunction remain unclear. While insulin secretion from pancreatic β-cells has long been known to be partly controlled by intracellular phospholipid signaling, very little is known about the role of phospholipids in glucagon secretion. Here we show that TMEM55A, a lipid phosphatase that dephosphorylates phosphatidylinositol-4,5-bisphosphate (PIP2) to phosphatidylinositol-5-phosphate (PI5P), regulates α-cell exocytosis and glucagon secretion. TMEM55A knockdown in both human and mouse α-cells reduces exocytosis at low glucose, and this is rescued by the direct reintroduction of PI5P. This does not occur through an effect on Ca^2+^ channel activity, but through a re-modelling of cortical F-actin dependent upon TMEM55A lipid phosphatase activity which occurs in response to oxidative stress. In summary, we reveal a novel pathway by which TMEM55A regulates α-cell exocytosis by manipulating intracellular PI5P level and the F-actin network.

## Introduction

Glucagon is secreted by α-cells of the pancreatic islets of Langerhans to stimulate glycogenolysis and gluconeogenesis, thereby increasing blood glucose [1]. In diabetes, hypersecretion of glucagon can contribute to postprandial hyperglycemia while an insufficient glucagon response to falling blood glucose levels can lead to hypoglycemia [1]. Both intrinsic and paracrine mechanisms control glucagon secretion, however in contrast to the exploration of detailed molecular mechanisms for insulin secretion from β-cells, the mechanisms regulating α-cell activity and glucagon secretion remain largely unclear and debatable [2]. Extrinsically, glucagon secretion is regulated by paracrine factors, such as insulin [3], somatostatin [4], adrenaline [5], and GIP [6]. Intrinsically, α-cell membrane depolarization initiates the opening of voltage-gated Na^+^ and P/Q-type Ca^2+^ channels, and a local increase of cytoplasmic Ca^2+^ which triggers the exocytosis of glucagon granules [1]. The access of those granules to the plasma membrane is also regulated by the cortical cytoskeleton [7]. Cortical F-actin is reported to act as a barrier to prevent access of insulin granules to the plasma membrane [8, 9], while also providing a path for insulin granule trafficking [10]. In contrast, studies on the relationship between cortical F-actin and glucagon granules in α-cells are sparse [11–13] and the up- or downstream regulatory pathway(s) remains poorly understood.

TMEM55A (encoded by the *PIP4P2* gene) was first identified as a phosphatase that catalyzes the hydrolysis of phosphatidylinositol-4,5-bisphosphate (PIP2) to phosphatidylinositol-5-phosphate (PI5P), with no enzymatic activity on other phosphoinositides [14]. It is expressed throughout the body, including in the pancreas [14]. Our published pancreas patch-seq data indicated that *PIP4P2* expression is positively correlated with α cell glucagon exocytosis, while negatively correlated with β cell insulin exocytosis [15]. We therefore explored whether and how TMEM55A regulates exocytosis in these cells. In the present study, we examined the role of TMEM55A in human and mouse primary α-cells, and the αTC1-9 cell line. We found that TMEM55A positively regulates α-cell exocytosis, by increasing intracellular PI5P levels to promote F-actin depolymerization via inhibition of the small G-protein RhoA. Oxidative stress acts upstream of TMEM55A/PI5P/F-active axis, resulting in increased glucagon exocytosis and glucagon hypersecretion.

## Results

### Expression of *PIP4P2*, encoding TMEM55A, regulates α-cell function

Our previous correlated electrophysiological and single-cell RNA-seq (patch-seq) studies of α-cells from human donors identified hundreds of transcripts correlated with α-cell function [15]. However, potential causative roles for most of these remain to be elucidated. Mining this data, a lipid phosphatase, TMEM55A (encoded by *PIP4P2*), stood out since it appeared to associate with single-cell function in α-cells. We confirmed the expression of TMEM55A in human α- and β-cells by immunostaining in biopsies from donors with and without T2D (Fig. 1A). Upon analysis of a large patch-seq dataset at www.humanislets.com [16], we found that the expression of *PIP4P2* is positively correlated with the α-cell size, exocytosis, Ca^2+^ currents and Na^+^ currents (Fig. 1B), while it does not correlate with most β-cell electrophysiological properties (Fig. 1C). We further compared the correlations in α-cells from donors with or without T2D (ND vs T2D) at 1 mM glucose and 5 mM glucose, and found that *PIP4P2* demonstrates a loss of correlation (or even opposite correlation) with α-cell electrophysiology parameters in T2D (Fig. 1D, E), indicating the function of TMEM55A might be dysregulated.

**Figure 1.**
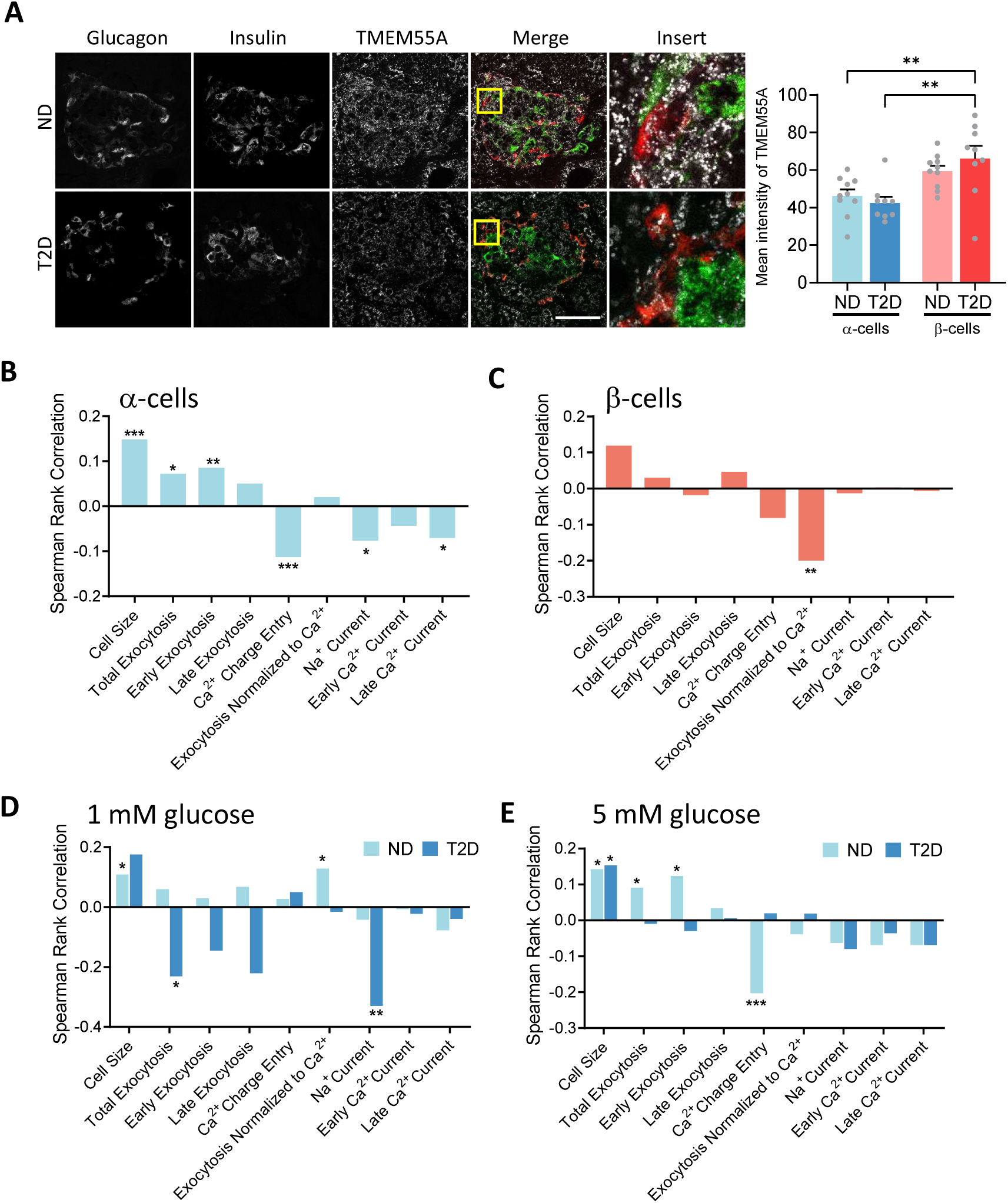
TMEM55A and islet cell function. **A.** Representative immunofluorescence images confirm TMEM55A expression in α- and β-cells of donors with no diabetes (ND) or type 2 diabetes (T2D). Scale bar, 50 µm. (n = 10 islets from 3 ND donors and 9 islets from 3 T2D donors) **B-C.** Correlation of *PIP4P2* transcript expression (encodes TMEM55A protein) with electrophysiological properties in human α-cells (B; n = 1114 cells from 45 donors) and β-cells (C; n = 229 cells from 37 donors). **D-E.** Correlation of *PIP4P2* transcript expression with electrophysiological properties in human α-cells at 1 mM glucose (D; n = 393 cells from 33 ND donors and 75 cells from 7 T2D donors) or 5 mM glucose (E; n = 524 cells from 43 ND donors and 211 cells from 11 T2D donors) from ND (light blue) or T2D (dark blue) donors. Data in panels B-E are from www.humanislets.com. * P < 0.05; ** P < 0.01; *** P < 0.001.

To better characterize the role of TMEM55A in islets, *PIP4P2* was knocked down by RNA interference (RNAi) in isolated human islet cells. Knockdown was confirmed at both mRNA (86% reduction) and protein (57% reduction) levels (Fig. 2A, B), and by single-cell immunofluorescence (Fig. 2C and Fig. S1A). Here we did not find any differences in the α-cell size or depolarization-induced Ca^2+^ entry upon the knockdown of *PIP4P2* but found that the α-cell exocytotic response (at 1 mM glucose) was dramatically decreased (Fig. 2D, E), consistent with the correlation analysis above. In β-cells, we found little effect following *PIP4P2* knockdown (Fig. S1 B, C). Since this single-cell exocytosis measurement can inform about mechanistic underpinnings of α-cell function, but does not measure secretion directly, we assessed glucagon secretion stimulated by KCl, low glucose (1 mM), and glucose-dependent insulinotropic polypeptide (GIP, 100 nM) with the amino acid alanine (10 mM) following *PIP4P2* knockdown. Consistently, we found that glucagon secretion from the si-*PIP4P2* transfection group was significantly reduced upon KCl or GIP and alanine stimulation compared with controls (Fig. 2F). These data together suggest the involvement of TMEM55A in the regulation of glucagon secretion.

**Figure 2.**
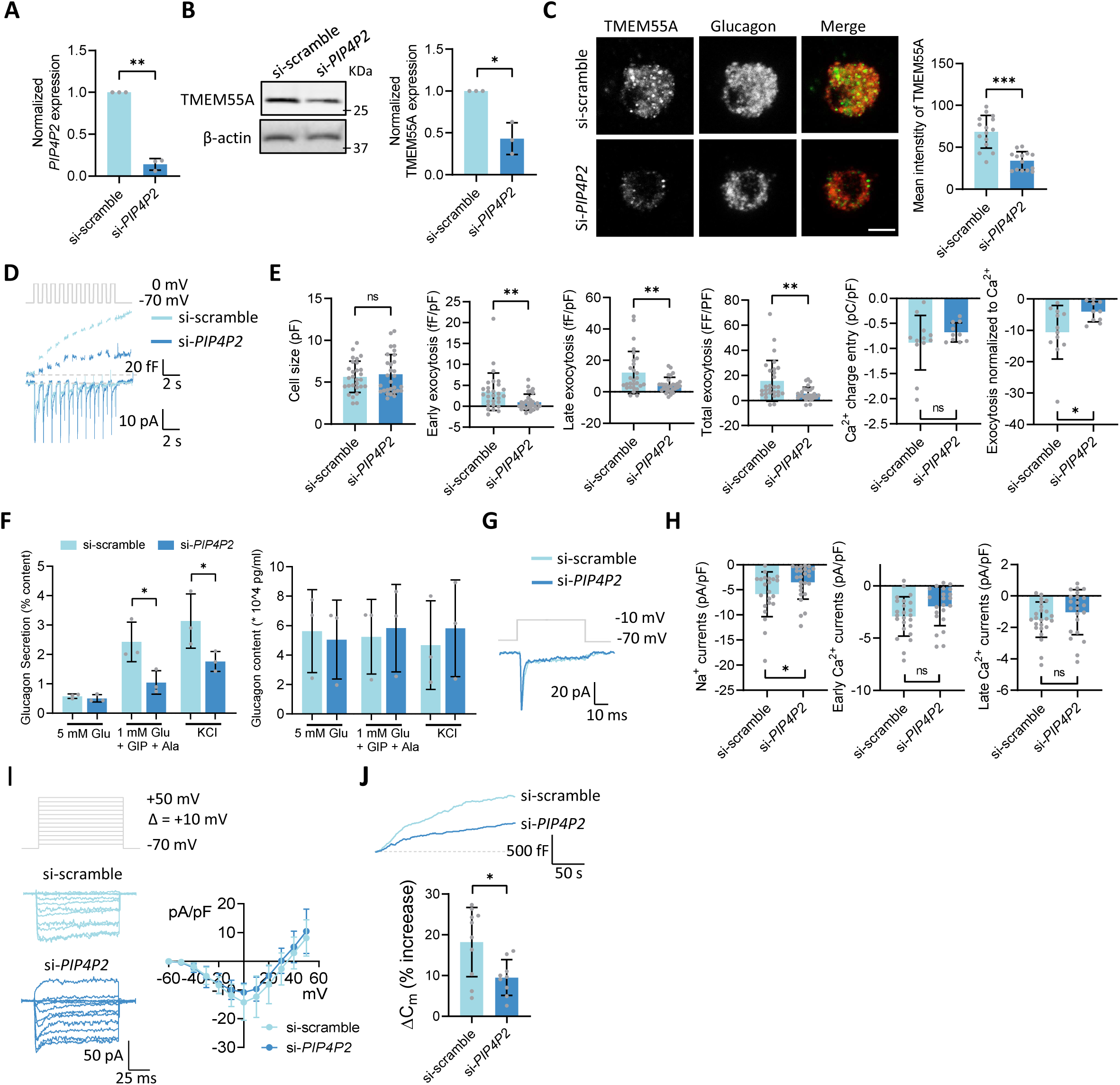
Knockdown of *PIP4P2* decreases α-cell exocytosis. **A.** qPCR analysis of *PIP4P2* mRNA expression from control (si-scramble) and *PIP4P2* knockdown (si-*PIP4P2*) human islet cells (n = 3 donors). **B.** Left panel: representative western blot of TMEM55A from control and *PIP4P2* knockdown human islet cells; Right panel: averaged blot intensities normalized to β-actin (n = 3 donors). **C.** Left panel: representative immunofluorescence images showing the TMEM55A from control and *PIP4P2* knockdown human α-cells. Scale bar, 5 µm. Positive glucagon staining is used to confirm the α-cell identity. Right panel: averaged intensities per cell (n = 15 and 14 cells from 3 donors). **D.** Representative capacitance and current traces induced by a train of 10 depolarizations from -70 mV to 0 mV (gray trace) from control and *PIP4P2* knockdown human α-cells. **E.** Averaged cell size (n = 28 and 29), early exocytosis (n =28 and 29), late exocytosis (n = 28 and 29), total exocytosis (n = 28 and 29), Ca^2+^ integral (n = 12 and 10) and normalized exocytosis to Ca^2+^ (n = 12 and 10), calculated as total exocytosis normalized to Ca^2+^ integral obtained from **D** (n = 5 donors). **F.** Left panel: glucagon secretion from dispersed control and *PIP4P2* knockdown human islets at basal (5 mM glucose) and stimulated conditions. Right panel: total glucagon content from left (n = 3 donors). **G.** Representative current traces induced by a depolymerization from -70 mV to - 10 mV (gray trace). **H.** Averaged Na^+^ currents (n =22 and 25), early Ca^2+^ currents (n = 24 and 22) and late Ca^2+^ currents (n = 24 and 22) from **G** (n = 5 donors). **I.** Left panel: representative current traces obtained using a voltage jump protocol (gray trace) from control and *PIP4P2* knockdown human α-cells. Right panel: averaged and normalized I-V curves obtained from left panel (n = 13 and 11 cells, from 3 donors). **J.** Upper panel: representative capacitance traces following 200 nM free Ca^2+^ infusion from control and *PIP4P2* knockdown human α-cells. Lower panel: averaged and normalized capacitance increase (ΔC_m_) after 200 s infusion obtained from left panel (n = 10 and 9 cells, from 3 donors). Data are presented as mean ± SD. Student’s t test (panels A-C, E, H, J), or one-way ANOVA and Holm-Sidak post-test (panel F). * P < 0.05; ** P < 0.01; *** P < 0.001; ns – not significant.

We also measured voltage-dependent Na^+^ and Ca^2+^ currents. The si-*PIP4P2* transfected α-cells showed decreased Na^+^ currents. However, we did not detect any significant changes in the Ca^2+^ currents (Fig. 2G, H). Consistent with the little effect on β-cell exocytotic response, no differences on the Na^+^ or Ca^2+^ currents were found upon *PIP4P2* knockdown in β-cells (Fig. S1 D, E). Since the intracellular Ca^2+^ inhibits voltage-dependent Ca^2+^ channels, we also used Ba^2+^ as a charge carrier and included the Na^+^ channel inhibitor tetrodotoxin (0.5 µM) in the bath to more closely interrogate Ca^2+^ channel activity. We still found no significant effect of *PIP4P2* knockdown on the Ca^2+^ channel-mediated currents (Fig. 2I), indicating that reduced Ca^2+^ channel activity is not responsible for the decreased α-cell exocytosis we observed above. Indeed, upon infusion of 200 nM free Ca^2+^ into the cell interior, exocytosis from the si-*PIP4P2* transfected α-cells was still lower than controls (Fig. 2J).

### A role for PIP2 and PI5P regulating α-cell exocytosis

TMEM55A was initially identified as a phosphatase that dephosphorylates PIP2 to PI5P [14]. As a signaling phospholipid, PIP2 plays critical roles in exocytosis [17, 18], while the physiological relevance of PI5P to exocytosis has never been explored. We therefore examined whether PIP2 or PI5P can regulate α-cell exocytosis. The intracellular dialysis of 1 µM diC_8_ PIP2 or PI5P, a water-soluble dioctanoyl analog, increased and decreased human α-cell exocytosis, respectively (Fig. 3 A, B), directionally consistent with what we would expect based on our TMEM55A knockdown studies. Indeed, PI5P infusion rescued the reduced exocytosis in si-*PIP4P2* transfected cells to levels comparable with the controls (Fig. 3C). Similar experiments could also be reproduced using mouse α-cells (Fig. 3 D, E). To better illustrate the role of PIP2 and PI5P on glucagon secretion, we treated mouse islets with 1 µM PIP2 or PI5P (which are membrane permeable [19, 20]) overnight and found that PIP2 has no obvious effect, while PI5P-treated islets demonstrate increased glucagon secretion (Fig. 3F). This suggests that the decreased glucagon secretion caused by *PIP4P2* knockdown might be due to the decreased intracellular PI5P level, rather than increased PIP2. Of note, neither PIP2 nor PI5P affects glucagon content (Fig. 3G), which is consistent with our knockdown study (Fig. 2F).

**Figure 3.**
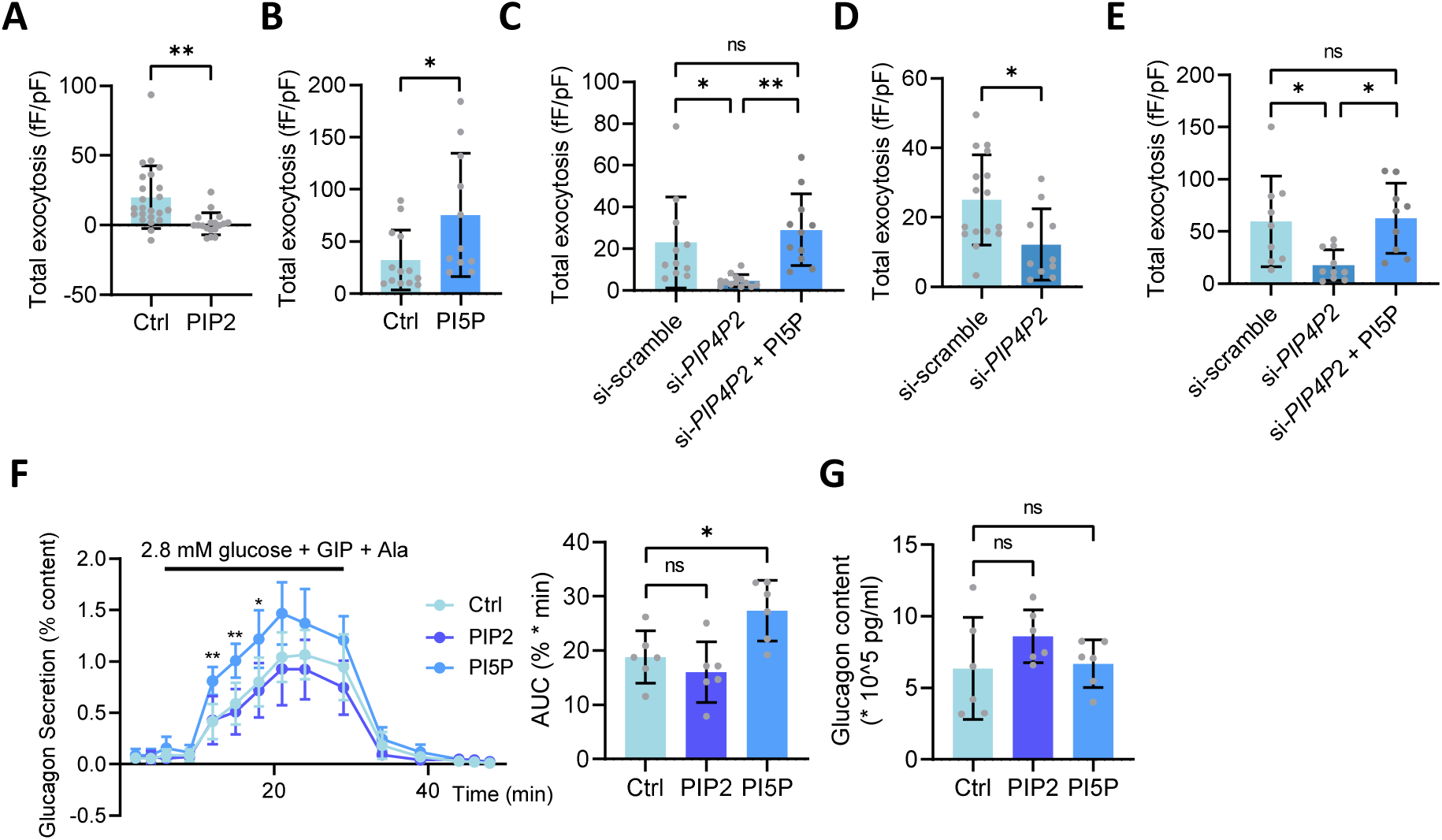
The effect of PIP2 and PI5P on glucagon secretion. **A.** Averaged total exocytosis with or without the infusion of 1 µM PIP2 in human α-cells (n = 23 and 18 cells, from 4 donors). **B.** Averaged total exocytosis with or without the infusion of 1 µM PI5P in human α-cells (n = 13 and 11 cells, from 3 donors). C. Averaged total exocytosis from control, *PIP4P2* knockdown, and *PIP4P2* knockdown with PI5P infusion human α-cells. (n = 11, 13 and 11 cells, from 3 donors). **D.** Averaged total exocytosis from control and *PIP4P2* knockdown mouse α-cells (n = 16 and 10 cells, from 3 mice). **E.** Averaged total exocytosis from control, *PIP4P2* knockdown, and *PIP4P2* knockdown with PI5P infusion mouse α-cells (n = 9, 10 and 9 cells, from 3 mice). **F.** Left panel: glucagon secretion induced by 2.8 mM glucose, GIP and Alaine from control mouse islets and islets with PIP2 or PI5P treatments. Right panel: AUC (area under the curve) obtained from left panel (n = 6 mice). **G.** Total glucagon content from F. Data are presented as mean ± SD. Student’s t test (panels A, B, D), Two-way ANOVA test with mixed-effects analysis (panel F) or one-way ANOVA followed by Holm-Sidak post-test (panels C, E, G). * P < 0.05; ** P < 0.01; ns – not significant.

### Phosphatase activity of TMEM55A

It has been suggested that TMEM55A does not, in fact, have lipid phosphatase activity [21]. To investigate this, and to confirm whether TMEM55A lipid phosphatase activity is involved in the regulation of α-cell function, we generated a wild-type GFP-tagged human TMEM55A (GFP-TMEM55A) and “phosphatase dead” mutant (GFP-TMEM55A C107S). Interestingly, human embryonic kidney (HEK) cells expressing GFP-TMEM55A demonstrated a more round-shape, compared with cells expressing GFP or GFP-TMEM55A C107S (Fig. S2 A, B), suggesting some activity of the WT enzyme influencing cell morphology. Moreover, we detected a large hydrolysis of PIP2 by GFP-TMEM55A, compared with GFP or GFP-TMEM55A C107S, which is even more robust compared with the positive control, the phosphatase enzyme SHIP2 (Fig. 4A). Since PI5P was reported as an oxidative stress-induced second messenger and it is increased in response to hydrogen peroxide (H₂O₂) in diverse cell lines [22], we examined whether TMEM55A activity could be modulated by cellular redox. Here we found that under reducing condition with 1 mM DTT, TMEM55A lipid phosphatase activity is inhibited while subsequent incubation with an oxidizer, 1 mM H_2_O_2_ activates the enzyme (Fig. 4B).

**Figure 4.**
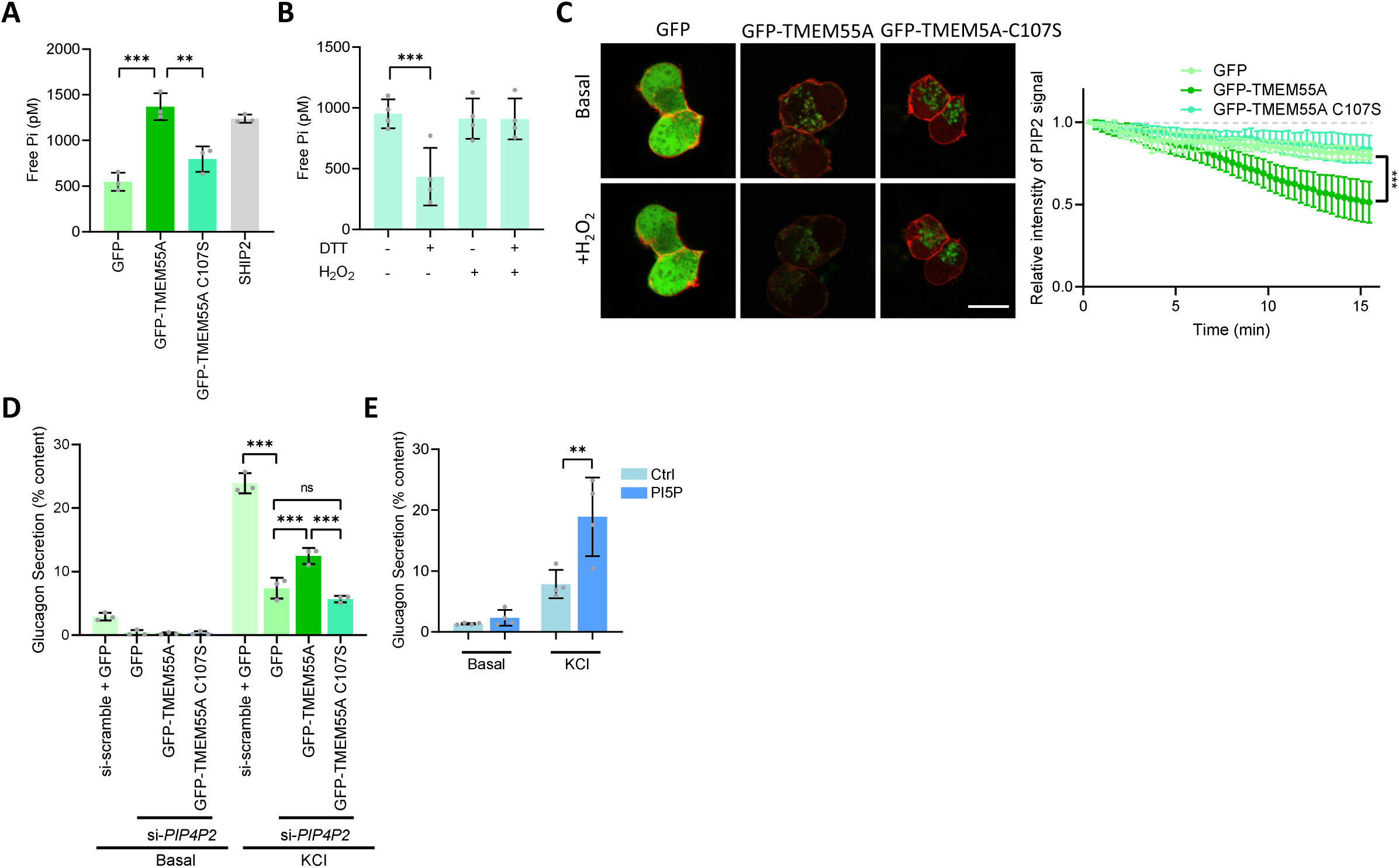
TMEM55A regulating glucagon secretion requires its phosphatase domain. **A.** Averaged phosphatase activity from GFP, GFP-TMEM55A, GFP-TMEM55A C107S and SHIP2 (n = 3). **B.** Averaged phosphatase activity of GFP-TMEM55A with the treatment of DTT, H_2_O_2_ or DTT and H_2_O_2_ (n = 4). **C.** Left panel: representative images of αTC1-9 cells transfected with RFP-PH-PLC (red) and GFP (green), GFP-TMEM55A, or GFP-TMEM55A C107 at the basal level and after treatment with H_2_O_2_ (1 mM) for 15 min. Scale bar, 10 µm. Right panel: quantification of the dynamic change of the PIP2 signal intensity in the αTC1-9 cells upon H_2_O_2_ stimulation from left (n = 8, 10 and 10 cells, from 3 independent experiments). **D.** Static glucagon secretion stimulated by KCl (55 mM) from αTC1-9 cells transfected with si-scramble + GFP, si-*PIP4P2* + GFP, si-*PIP4P2* + GFP-TMEM55A, si-*PIP4P2* + GFP-TMEM55A C107S (n = 3). **E.** Static glucagon secretion stimulated by KCl (55 mM) from αTC1-9 cells with or without the treatment of PI5P (n = 4). Data are presented as mean ± SD. Two-way ANOVA test with mixed-effects analysis (panel C) or one-way ANOVA followed by Holm-Sidak post-test (panels A , B, D, E). * P < 0.05; ** P < 0.01; *** P < 0.001; ns – not significant.

Next, we tested the H_2_O_2_ sensitivity of TMEM55A *in situ*. Because there is no reliable PI5P sensor, we overexpressed GFP-TMEM55A and PIP2 probe, mCherry-PH-PLC [23] at the same time. Then, we monitored the dynamic PIP2 changes following treatment with 1 mM H_2_O_2_ using live cell imaging. As reported before [24], external application of H_2_O_2_ induced PIP2 hydrolysis. Interestingly, while treatment with H_2_O_2_ activated PIP2 hydrolysis in cells expressing wild-type GFP-TMEM55A, this effect was lost in cells expressing GFP-TMEM55A C107S (Fig. 4C and supplementary video 1-3), indicating that H_2_O_2_ induced PIP2 hydrolysis depends on the phosphatase activity of TMEM55A.

We then knocked down native TMEM55A in αTC1-9 cells. Like in primary α-cells, knocking down TMEM55A decreased glucagon secretion from αTC1-9 cells (Fig. 4D). When we overexpressed GFP-TMEM55A and GFP-TMEM55A C107S, we found that only the wild-type GFP-TMEM55A showed a partially recovered glucagon secretion following TMEM55A knockdown (Fig. 4E). This indicates that TMEM55A control of glucagon secretion depends on its phosphatase activity. Moreover, the product of TMEM55A, PI5P significantly increases glucagon secretion in αTC1-9 cells (Fig. 4F). Taken together, these data suggest that the regulation of TMEM55A on glucagon secretion depends on its lipid phosphatase activity.

### RhoA is the downstream signaling molecule of PI5P

PI5P is the least characterized phosphoinositide and its functions remain elusive [25]. It was reported that PI5P might regulate actin dynamics in cell migration [26], and intracellular or extracellular application of PI5P in different cell lines causes F-actin stress fiber breakdown via activation of Rac1[20, 27]. To address the downstream signaling effect of PI5P in α-cells, we monitored the levels of active Rac1 together with two other major GTPases, Cdc42 and RhoA in αTC1-9 cells following treatment with PI5P. We found that PI5P leads to strong inactivation of RhoA, with no obvious effects on Rac1 or Cdc42 (Fig. 5A). Although this contradicts observations in mouse embryonic fibroblast (MEF) cell lines [20], it is in line with the role of RhoA identified in α-cells [12]. Similar experiments could also be reproduced in human islets (Fig. 5B). Expression of GFP-TMEM55A in αTC1-9 cells also resulted in the inactivation of RhoA, while GFP or GFP-TMEM55A C107S had no effect (Fig. 5C). These observations suggested that RhoA is a downstream effector of PI5P and TMEM55A in α-cells.

**Figure 5.**
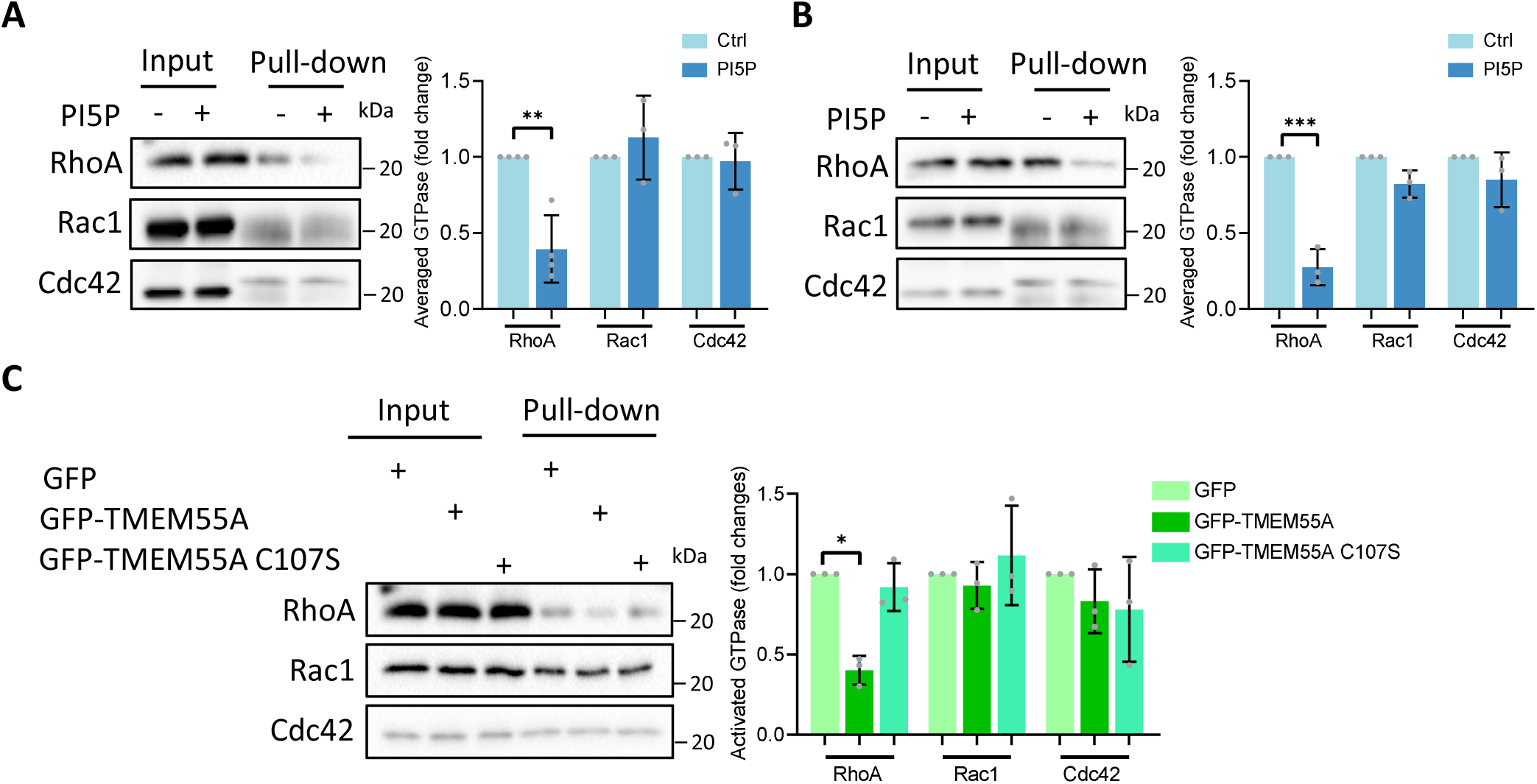
PI5P and TMEM55A inactivate RhoA, while have no effects on Rac1 and Cdc42. **A.** Left panel: pull-down assays showing the levels of active forms of Rac1, Cdc42 and RhoA in αTC1-9 cells treated with or without PI5P (10 µM) for 1 h. Right panel: averaged and normalized active form of GTPases from left (N = 4, 3, 3). **B.** Left panel: pull-down assays showing the levels of active form of Rac1, Cdc42 and RhoA for human islets treated with or without PI5P (1 μM) overnight. Right panel: averaged and normalized active form of GTPases from left (n = 3, 3, 3) **C.** Left panel: pull-down assays showing the levels of active form of Rac1, Cdc42 and RhoA for GFP, GFP-TMEM55A or GFP-TMEM55A C107S transfected αTC1-9 cells treated with or without PI5P (10 µM) for 1 h. Right panel: averaged and normalized active form of GTPases from left (n = 3, 3, 3). Data are presented as mean ± SD. Student’s test (panels A, B) or one-way ANOVA followed by Holm-Sidak post-test (panels C). * P < 0.05; ** P < 0.01; *** P < 0.001.

### PI5P regulates α-cell F-actin remodeling

F-actin depolymerization is mainly governed by Rho-family small GTPases [28]. After validation that RhoA is downstream of TMEM55A in α-cells, we wanted to visualize whether PI5P could disrupt the F-actin in α-cells to enhance glucagon secretion. We first confirmed the relationship between F-actin and α-cell exocytosis. Human α-cells treated with 10 µM latrunculin B or jasplakinolide for 1 h to disrupt and enhance actin polymerization, demonstrated increased and decreased exocytotic responses, respectively, directionally consistent with the effects of TMEM55A and PI5P (Fig. 6A). Then we examined the distribution of F-actin by phalloidin staining using confocal microscopy following PI5P treatment. Consistent with previous studies on MEF and HeLa cell lines [20, 27], PI5P significantly decreased the F-actin intensity in dispersed human and mouse primary α-cells (Fig. 6A, B), in whole islets (Fig. 6C, D), and in αTC1-9 cells (Fig. 6E). Using live-imaging of F-tractin-mCherry, a probe for monitoring global F-actin [29], we found that PI5P disrupted F-actin in a time-dependent manner (Fig. 6F and supplementary video 4-5). Since TMEM55A increases intracellular PI5P, we overexpressed GFP-TMEM55A in αTC1-9 cells and found that αTC1-9 cells overexpressing GFP-TMEM55A demonstrated a low intensity of F-actin compared with cells expressing GFP-TMEM55A C107S or GFP alone (Fig. 6G). Conversely, when we knocked down *PIP4P2* in human α-cells, we detected a significant increase of F-actin (Fig. 6 H), which explains the decreased exocytosis in our functional studies (Fig. 2E). Taken together, these data strongly indicated the involvement of TMEM55A and PI5P in F-actin depolymerization.

**Figure 6.**
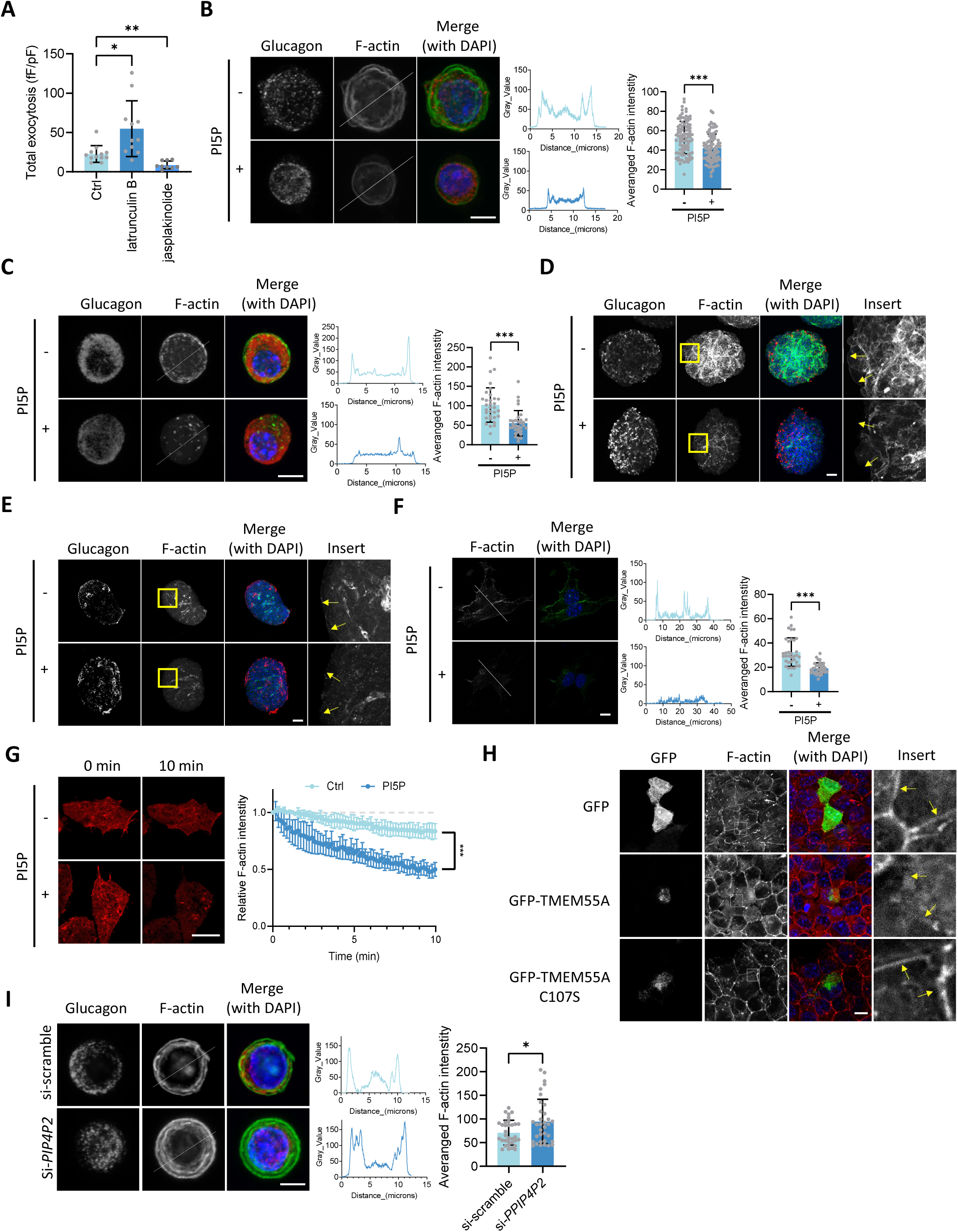
F-actin depolymerization is involved in TMEM55A’s regulation on glucagon secretion. **A.** Averaged total exocytosis with or without the incubation of 10 µM latrunculin B or jasplakinolide for 1 h in human α-cells (n = 12, 11 and 9, from 3 donors). **B, C and F.** Left panel: representative immunofluorescence images of human (B) and mouse (C) α-cells and αTC1-9 cells (F) with or without PI5P treatment for 1 h. F-actin was visualized by staining with AlexaFluor 647-phalloidin. Positive glucagon staining is used to confirm the α-cell identify from dispersed human or mouse islets. Scale bar, 5 µm. Right panel: F-actin fluorescence intensity line profile analysis from left and averaged F-actin intensity per cell (n = 96 and 85 cells, from 5 donors; n = 32 and 32 cells, from 3 mice; n = 34 and 34 cells, from 3 independent experiments). **D, E.** Representative immunofluorescence images of human (D) and mouse (E) whole islets with or without PI5P treatment overnight from three independent experiments. Yellow arrows indicate the disruption of F-actin. Scale bar, 50 µm **G.** Left panel: representative images of αTC1-9 cells transfected F-tractin-mCherry at the basal level and after treatment with PI5P (10 µM) for 10 min. Scale bar, 10 µm. Right panel: quantification of the dynamic change of the F-actin signal intensity in the αTC1-9 cells upon PI5P treatment from left (n = 6 and 7 cells, from 3 independent experiments). **H.** Representative immunofluorescence images of αTC1-9 cells transfected with GFP, GFP-TMEM55A or GFP-TMEM55A C107S from three independent experiments. Yellow arrows indicate the disruption of F-actin. Scale bar, 10 µm **I.** Left panel: representative immunofluorescence images showing the F-actin level from control and *PIP4P2* knockdown human α-cells. F-actin was visualized by staining with AlexaFluor 647-phalloidin. Positive glucagon staining is used to confirm the α-cell identity. Scale bar, 5 µm. Right panel: F-actin fluorescence intensity line profile analysis from left and averaged F-actin intensity per cell (n = 33 and 34 cells, from 3 donors). Data are presented as mean ± SD. Student’s t test (panels B, C, F, I), one-way ANOVA followed by Dunnett’s T3 multiple comparison test (panel A), or two-way ANOVA test with mixed-effects analysis (panel G). * P < 0.05; ** P < 0.01; *** P < 0.001.

## Discussion

While T2D results from a complex interplay between insulin resistance and insulin secretion [30, 31] and T1D results from insulin insufficiency due to autoimmune description of β-cells [32], both forms of diabetes are associated with disrupted glucagon secretion from islet α-cells [33]. However, the underlying molecular mechanisms that regulate glucagon secretion in α-cells remain unclear. Here we identified TMEM55A as a correlate of α-cell function through bioinformatic analysis of single-cell function and transcript expression. Combined with mechanistic studies on human and mouse primary α-cells and αTC1-9 cell lines, we describe a glucagon regulation mechanism by TMEM55A and PI5P-mediated F-actin depolymerization. We further provide evidence that oxidative stress can modulate TMEM55A activity, and that RhoA acts as a downstream effector of this lipid phosphatase, which is summarized in Fig. 7.

**Figure 7.**
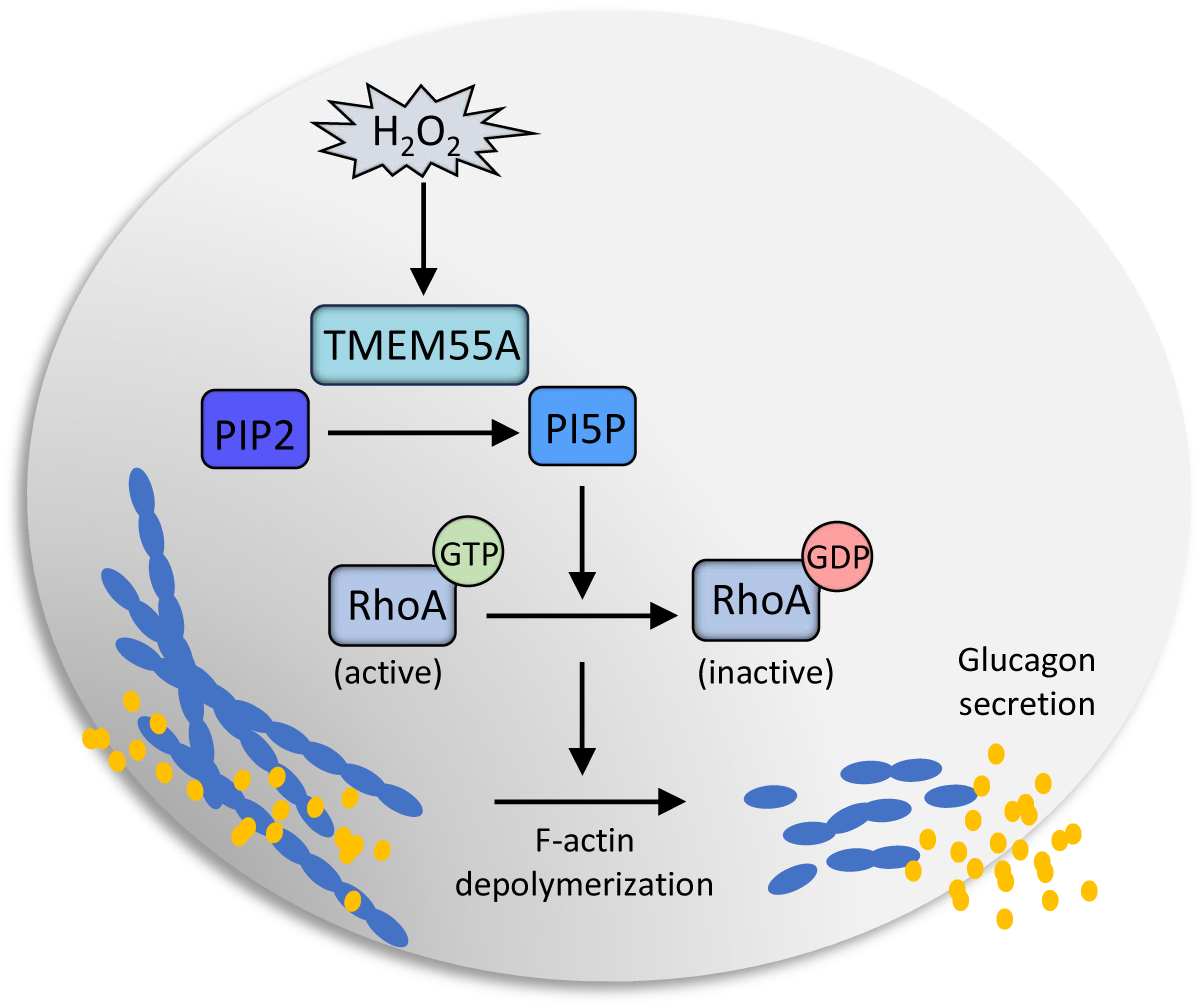
Schematic model showing how TMEM55A regulates glucagon secretion. In this model, upstream oxidative signals, such as H_2_O_2_, activates TMEM55A. TMEM55A positively regulates glucagon secretion by dephosphorylating PIP2 to PI5P and induces RhoA dependent F-actin depolymerization in pancreatic α-cells.

TMEM55A was initially identified, along with TMEM55B, as an enzyme that catalyzes PIP2 to PI5P [14]. However, its phosphatase activity has been challenged by recent reports [21, 34], which found that neither recombinant TMEM55A nor TMEM55B catalytic domain demonstrates any detectable phosphatase activity when expressed in *E.coli*. We assumed that the lack of activity could be due to the lack of transmembrane or post-translational modifications [35, 36]. We therefore expressed full-length TMEM55A in mammalian cells and performed the *in vitro* phosphatase assay on beads, with a loss-of function mutant serving as a negative control and could easily detect robust phosphatase activity. This suggests that modifications or binding partners conferred in mammalian cells are essential for TMEM55A phosphatase activity. Furthermore, live-cell imaging using PIP2 probes also strongly indicates the *in situ* function of TMEM55A which is under the control of redox state. Consistent with this, TMEM55A knockdown in mouse macrophage cell lines leads to the accumulation of PIP2 and a decrease of PI5P [37].

As a dense layer underneath the plasma membrane, F-actin is dynamically remodeled following intra- and extra-cellular signaling [38, 39]. Glucose-stimulated F-actin reorganization is critical for insulin secretion in β-cells [8, 9], although this has been challenged recently [40]. We showed previously that impaired regulation of β-cell actin remodeling by atypical phosphatidyl inositol 3-OH kinases (PI3Ks) contributes to dysfunction in T2D [41]. The PI3K signaling pathway also regulates insulin secretion by recruiting insulin granules, modifying cAMP, and interleukin-1 signaling [42–44]. However, the role of phosphoinositide signaling pathways and F-actin remodeling on glucagon secretion has not been explored. Here, we found that F-actin mainly acts as a negative regulator of glucagon secretion, which agrees with other studies [11, 12]. Further, we identified that the inhibition of RhoA acts as a downstream signaling event of PI5P. This is in contrast to MEF cells where Rac1 and Cdc42, but not RhoA activity, were found to be altered following PI5P treatment [20]. Currently, we lack an understanding of the mechanism by which PI5P regulates RhoA. Viaud *et al*. demonstrated that PI5P physically interacted with the exchange factor Tiam1, leading to the restricted Rac1 activation [20]. Numerous PI5P-interacting proteins were identified using yeast proteome microarrays combined with flow cytometry [45]. It is possible that some adapter proteins are involved in PI5P’s action on RhoA, which requires additional study.

We previously demonstrated that glucose suppresses human and mouse α-cell exocytosis, while non-metabolizable glucose analog 2-DG does not, due to the acute glucose treatment suppressing P/Q Ca^2+^ channel activity via complex I-dependent production of reactive oxygen species/H_2_O_2_ [15]. However, glucose could also increase α-cell exocytosis because of different culture conditions and the missing islet paracrine signaling [15, 46]. Even a U-shaped response of isolated human α-cells to glucose was reported [47], indicating a potential dual effect of oxidative signaling on glucagon secretion. H_2_O_2_ was reported to increase intracellular PI5P levels in human osteosarcoma cells, independently of mTOR, PDK1, PKB, ERK, p38 or PIKfyve signaling [22]. Here we found that H_2_O_2_ could also increase PI5P (indirectly monitored as a reduction in PIP2) in α-cells by activating TMEM55A. Considering in Hela cells that the type I phosphatidylinositol phosphate 5-kinase (PIP5Kbeta) is involved in the response to H_2_O_2_ [24], it is possible that diverse PIP2 kinases and phosphatases are critical in α-cell function.

Our correlational studies showed that α-cells with high *PIP4P2* expression in T2D demonstrate lower exocytosis. Moreover, knocking down *PIP4P2* in α-cells from T2D donors shows an unexpectedly higher exocytosis (Fig. S3A), suggesting that besides RhoA-dependent F-actin depolymerization, PI5P might have additional roles in T2D α-cells. We also tried to measure the total glucagon exocytosis from mouse *PIP4P2* knockdown α-cells following 14-weeks high fat diet (HFD) to mimic the T2D condition and we identified that TMEM55A still acts as a positive regulator for glucagon secretion under this condition (Fig. S3B), suggesting the differences between human and mouse model, which has been described before [15]. Compared with cytoplasmic PI5P, nuclear PI5P was also reported to regulate gene expression and apoptosis during stress response [48, 49]. It’s likely that a potential excessive PI5P plays a dominant role in the nucleus during diabetes. In both human and mouse α-cells, our data indicates that TMEM55A positively regulates glucagon secretion in the absence of diabetes.

In summary, our current study highlights the importance of TMEM55A in regulating α-cell exocytosis and glucagon secretion. TMEM55A is activated in response to oxidative stress, such as H_2_O_2_, to dephosphorylate PIP2 to PI5P, which inhibits the activation of RhoA. Inhibition of RhoA will depolymerize F-actin and promote glucagon exocytosis. This TMEM55A/PI5P/F-actin regulation axis is critical for glucagon secretion.

## Methods

### Plasmids

The cDNA of human *PIP4P2* was prepared by PCR using the commercial TMEM55A ORF clone in a pcDNA 3.1 (Genscript; SC1200) as a template. The mammalian expression vector of GFP tagged human TMEM55A (GFP-TMEM55A) was generated by inserting the *PIP4P2* cDNA into PEGFPC1 plasmid that was obtained from Dr. Kiyomi Nigorikawa (Hiroshima University, Japan). Loss of function mutant (GFP-TMEM55A C107S) was generated using Q5 High-Fidelity 2X Master Mix (New England Biolabs, NEB) and verified by sequencing. Primers used are in Table S1. F-tractin-mCherry was purchased from Addgene (155218). RFP-PH-PLC was provided by Dr. Todd Alexander (University of Alberta, Canada).

### siRNA constructs and quantitative PCR

Human and mouse TMEM55A and scrambled siRNA were purchased from Horizon Discovery Ltd (Cambridge, UK; E-013808-00-0005 and E-059670-00-0005). The FAM-labeled Negative Control siRNA was from Thermo Fisher Scientific (Ottawa, ON, Canada; AM4620). These were transfected in dissociated mouse, human islet cells or αTC1-9 cells using Lipofectamine™ RNAiMAX transfection reagent (Thermo Fisher Scientific; 13778075) according to the manufacturer’s protocol. Quantitative PCR (qPCR) was performed as previously described [50], RNA from the corresponding cells was extracted 72 h post-transfection using TRIzol reagent (Life Technologies, Burlington, ON), and the cDNA was synthesized using OneScript® Plus cDNA Synthesis Kit (ABM, Richmond, BC, Canada). Real-time PCR was carried out on a 7900HT Fast Real-Time PCR system using PowerUp™ SYBR™ Green Master Mix (Thermo Fisher Scientific; A25742). Primers used is in Table S1.

### Cell culture and transfection

αTC1-9 cells (a gift from Dr. Peter Light, University of Alberta, Canada) were cultured in low glucose DMEM media supplemented with 10% FBS, 0.02% BSA, 15mM HEPES, 100 µM non-essential amino acids, 100 U/mL penicillin/streptomycin. The cells were kept at 37 °C with 10% CO_2_. Human Embryonic Kidney (HEK) 293 cells (a gift from Dr. Harley Kurata, University of Alberta, Canada) were cultured in high glucose DMEM media supplemented with 10% FBS and 1% penicillin/streptomycin. The cells were kept at 37 °C with 5% CO_2_. Transfection of cDNAs was performed using Lipofectamine 3000 (Invitrogen; L3000008) according to the manufacturer’s protocol.

### Human islets

Human islets were from our Alberta Diabetes Institute IsletCore (www.isletcore.ca) [51], the Clinical Islet Transplant Program at the University of Alberta, or the Human Pancreas Analysis Program [52]. Human islets were cultured in DMEM media supplemented with 10 % FBS and 1% penicillin/streptomycin. The islets were kept at 37 °C with 5% CO_2_. This work was approved by the Human Research Ethics Board at the University of Alberta (Pro00013094; Pro00001754) and all families of organ donors provided written informed consent.

### Mouse islets

Mouse islets were isolated from male C57BL/6NCrl mice (Charles River Laboratories) fed with standard chow diet at 10-12 weeks of age. Mouse islets were cultured in RPMI-1640 with 11.1 mM glucose, 10 % FBS, 1% penicillin/streptomycin. The present mouse islet study was approved by the Animal Care and use Committee at the University of Alberta (AUP00000291).

### Patch-clamp recordings

Patch-clamp recordings were performed as described previously [15]. Hand-picked islets were dissociated into single cells using enzyme-free cell dissociation buffer (Gibco, 13150016) and cultured at 37°C with 5% CO_2_ for 1-3 days. For depolarization-stimulated exocytosis and voltage-gated Na^+^ and Ca^2+^ current measurements, media was changed to a bath solution containing (in mM): 118 NaCl, 20 Tetraethylammonium-Cl, 5.6 KCl, 1.2 MgCl_2_, 2.6 CaCl_2_, 5 HEPES, and 1 glucose (pH 7.4 with NaOH) in a heated chamber (37 °C). The recording pipettes with a resistance of 4-7 MΩ were fire polished, coated with Sylgard and filled with the following intracellular solution (in mM): 125 Csglutamate, 10 CsCl, 10 NaCl, 1 MgCl_2_, 0.05 EGTA, 5 HEPES, 0.1 cAMP, and 3 MgATP (pH 7.15 with CsOH). For assessment of Ca^2+^ channel activity using Ba^2+^ as a charge carrier, the bath solution was composed of (in mM): 100 NaCl, 20 BaCl_2_, 5 CsCl, 1 MgCl_2_, 10 HEPES, with 0.2 μM tetrodotoxin, 1 glucose (pH 7.4 with NaOH); and the intracellular solution was composed of (in mM): 140 Cs-glutamate, 1 MgCl_2_, 20 TEA, 5 EGTA, 20 HEPES, 3 Mg-ATP (pH 7.15 with CsOH). Electrophysiological data were recorded using a HEKA EPC10 amplifier and PatchMaster Software (HEKA Instruments Inc, Lambrecht/Pfalz, Germany) within 5 min of break-in. Patched cells were marked on the dish and subjected to immunostaining for insulin and glucagon for cell type identification.

### In beads phosphatase assay

HEK 293 cells were transfected with either GFP vector or GFP tagged-TMEM55A WT or C107S mutant using Lipofectamine 3000 (Invitrogen) according to the manufacturer’s protocol. Those cells were solubilized in ice-cold CelLytic-M lysis buffer (Sigma-Aldrich; C2978) supplemented with protease inhibitor cocktail (Sigma-Aldrich; 539137). Supernatants were collected after centrifugation at 13,200 rpm for 15 min, followed by incubation with GFP-Trap Agarose (Chromotek) at 4 °C. The mixture was incubated at 4 °C for 1 h with rotation end-over-end. Agarose beads were then washed three times using washing buffer (in mM): 10 Tris-Cl pH 7.5, 150 NaCl, 0.5 EDTA, and 0.05 % NP40 substitute. Bead complexes were resuspended with 50 μL reaction buffer (in mM): 25 Tris-Cl pH 7.4, 140 NaCl, 2.7 KCl, with or without adding 1 mM DTT or H_2_O_2_. Commercially purchased purified SHIP2 (Cedarlane) was mixed with the reaction buffer as a positive control. The reactions were processed immediately by adding diC_8_-PtdIns-4,5-P2 (PIP2) (Echelon Biosciences) at a final concentration of 1 mM with incubation at 37 °C for 1 h. Supernatants were collected after centrifugation at 3,000 rpm for 15 min. 100 μL Malachite Green Solution (Echelon Biosciences) was mixed with 25 μL sample or phosphate standard solution (0-2000 pM) in a well of the 96-well polystyrene microplate. After 20 mins incubation at room temperature, absorbance was read at 620 nm. The standard curve was prepared for every repeated experiment.

### GTPase activation assay

The activity of Rac1/Cdc42/RhoA was measured by RhoA/Rac1/Cdc42 Activation Assay Combo Biochem Kit (Cytoskeleton Inc, BK030) following a modified manufacturer protocol. Briefly, islets or αTC1-9 cells were lysed in ice-cold lysis buffer (in mM): 50 Tris pH 7.5, 10 MgCl_2_, 500 NaCl, and 2% igepal supplemented with protease inhibitor cocktail (Sigma-Aldrich). Supernatants were collected after centrifugation at 13,200 rpm for 15 min, followed by incubation with rhotekin-RBD (for RhoA activation assay) or PAK-PBD beads (for Rac1 and Cdc42 activation assays) at 4 °C overnight. Beads were pelleted by centrifugation at 3000 rpm at 4 °C for 1 min. Beads were then washed three times using washing buffer (in mM): 25 Tris pH 7.5, 30 MgCl_2_, 40 NaCl and eluted by SDS loading buffer. Precipitated active forms of GTPases were subjected to western blot analysis.

### Immunofluorescence

Cells on the coverslips or islets were washed with PBS and fixed in 4% paraformaldehyde for 15 min at room temperature, then permeabilized with 0.1% Triton X-100 for 10 min. After permeabilization, cells or islets were blocked in PBS plus 3% BSA for 30 min at room temperature and then incubated overnight with indicated primary antibodies in the cold room, followed by incubation with corresponding secondary antibodies or together with Phalloidin-iFluor 647 Reagents (Abcam, Cambridge, UK) for 1 h at room temperature. Cells or islets were then mounted on the glass slides and examined using Leica TCS SP5 confocal laser scanning microscope (Cell Imaging Facility, Faculty of Medicine and Dentistry, University of Alberta).

### Live cell imaging

αTC1-9 cells were seeded into the Cellvis 35 mm glass bottom dish with 14 mm micro-well #1.5 gridded (interior) cover glass (Thermo Fisher Scientific) and transfected with F-tractin-mCherry or GFP-PH-PLC to visualize F-actin and PIP2, respectively. Cells were pre-incubated with DMEM without any supplements 24 h before imaging. Dynamic signaling following the stimulation was captured and recorded every 10 or 20 s for 10 or 15 min using Leica Stellaris 8 confocal microscope equipped with a live-cell chamber (Cell Imaging Facility, Faculty of Medicine and Dentistry, University of Alberta). The fluorescence signal intensity of F-actin or PIP2 was quantified by ImageJ.

### Glucagon secretion

For static glucagon secretion measured with dispersed human islets, 50 siRNA transfected islets per group were pre-incubated for 30 min in KRB buffer containing (in mM): 140 NaCl, 3.6 KCl, 2.6 CaCl_2_, 0.5 NaH_2_PO_4_, 0.5 MgSO_4_, 5 HEPES, 2 NaHCO_3_ and 0.5 mg/ml BSA supplied with 5 mM glucose. Then they were stimulated with or without glucose (1 mM), GIP (100 nM) and the amino acid alanine (10 mM), or KCl (20 mM). For static glucagon secretion measured with αTC1-9 cells, cells plated on 12-well plates were serum starved with DMEM without any supplements the day before the secretion assay. Cells were washed twice with PBS and then incubated with fresh DMEM for 1 h. KCl (55 mM) was used to stimulate the glucagon secretion. For some experiments, PI5P was added to the fresh DMEM for 1 h incubation. For dynamic glucagon responses, 30 islets from each group were pre-incubated for 30 min in KRB buffer containing (in mM): 140 NaCl, 3.6 KCl, 2.6 CaCl_2_, 0.5 NaH_2_PO_4_, 0.5 MgSO_4_, 5 HEPES, 2 NaHCO_3_ and 0.5 mg/ml essentially fatty acid free BSA. And then they were perifused in the same buffer with the changes of indicated glucose concentration and the addition of GIP (100 nM) and the amino acid alanine (10 mM). Total glucagon content was obtained by lysing the cells with acid ethanol supplemented with protease inhibitor cocktail. All the samples were collected and stored at -20 °C for assay of glucagon with either Lumit Glucagon kit (Promega; W8020) or glucagon ELISA kit (MSD; K1515YK-2).

### Correlation analysis of *TMEM55A* expression against electrophysiology in patch-seq data

Raw sequencing reads are available at the NCBI Gene Expression Omnibus under the following accession numbers: GSE270484, GSE124742 and GSE164875, and further available at www.humanislets.com with associated electrophysiology and donor characteristic. Correlations were performed using Log_2_[CPM+1] values for *PIP4P2* expression. Electrophysiology was corrected for outliers by only including values with a z-score between 3 and -3 for each respective correlation. For total, early, and late capacitance, negative values were considered noise and set to 0 to reduce their effects on correlations, as they are irrelevant in contributing to functional responses for exocytosis [53]. Data import, correlations with statistics, and export were performed in Python (v 3.7.11) using numpy (v 1.21.6), pandas (v 1.3.2), and (scipy 1.7.3) packages.

## Author contributions

Conceptualization, X.L. and P.E.M.; Investigation, X.L., T.dS., A.F.S., S.D., N.S., K.S.; Supervision, P.E.M.; Writing, X.L., P.E.M.

## Acknowledgements

The University of Alberta is situated on Treaty 6 territory, the traditional land of First Nations and Métis people. We thank Give Live Alberta and Trillium Gift of Life Network (TGLN) for their work in procuring human donor pancreas for research. We also thank James Lyon and Nancy Smith (Alberta) for their efforts in human islet isolation, and Kiera Smith (Alberta) from Cell Imaging Core - Katz Group Centre for her assistance on live-cell imaging experiments. We especially thank the organ donors and their families for their kind gift supporting diabetes research.

This work was supported by a Foundation Grant (FS 148451) and Project Grant (PS 186226) to PEM from the Canadian Institutes of Health Research. We also acknowledge the Human Pancreas Analysis Program (HPAP-RRID:SCR_016202) (https://hpap.pmacs.upenn.edu), a Human Islet Research Network (RRID:SCR_014393) consortium (UC4-DK-112217, U01-DK-123594, UC4-DK-112232, and U01-DK-123716). This work includes data and/or analyses from HumanIslets.com funded by the Canadian Institutes of Health Research, JDRF Canada, and Diabetes Canada (5-SRA-2021-1149-S-B/TG 179092). XL was supported in part by fellowships from the Canadian Islet Research and Training Network NSERC-CREATE program and from Alberta Innovates – Health Solutions. PEM holds the Tier 1 Canada Research Chair in Islet Biology.

