## Supplementary information for "TMEM55A-mediated PI5P signaling regulates α-cell actin depolymerization and glucagon secretion"

Number of supplemental figures: 4

Number of supplemental tables: 2

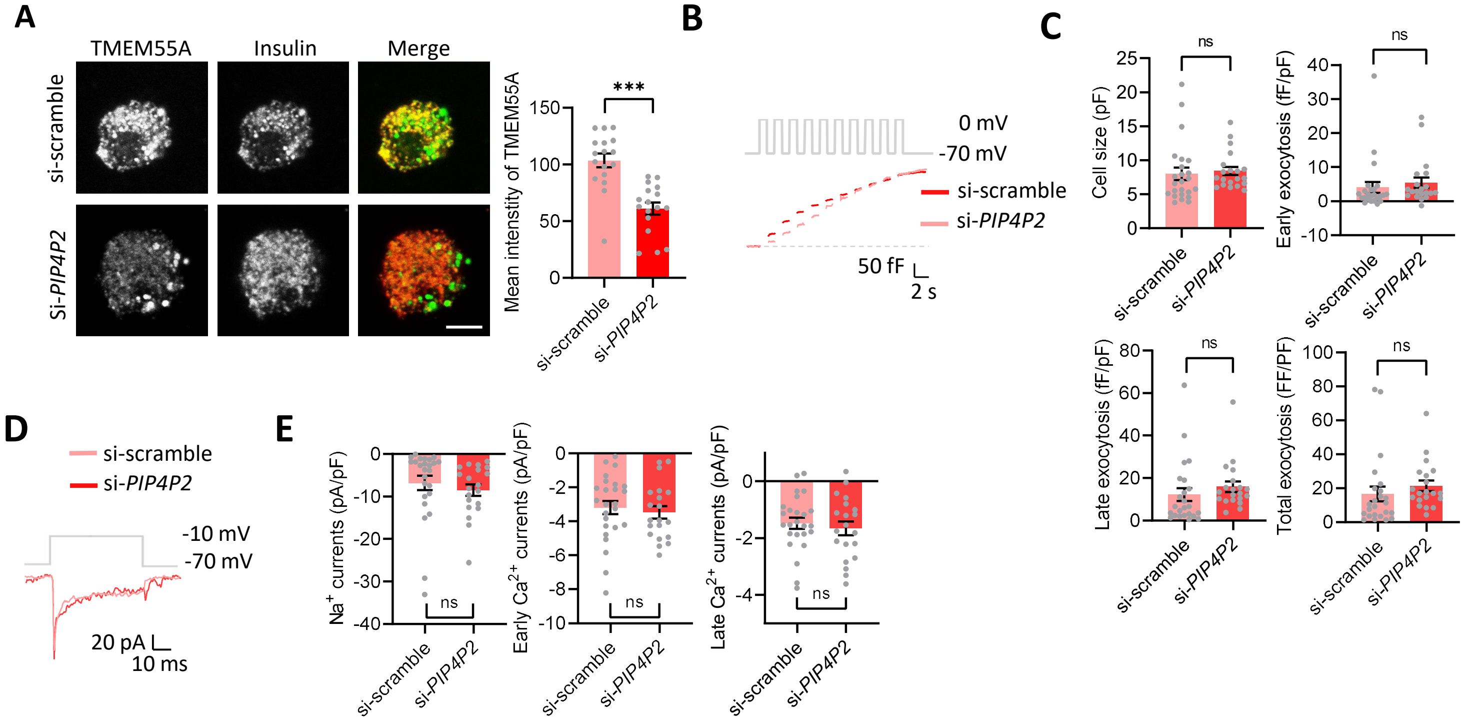


**Figure S1. Knockdown of TMEM55A does not affect β-cell exocytosis. A.** Left panel: representative immunofluorescence images showing the TMEM55A level from control and *PIP4P2* knockdown human β-cells. Scale bar, 5 µm. Positive insulin staining is used to confirm the β-cell identify. Right panel: averaged intensities per cell (n = 17 and 17 cells from 3 donors). **B.** Representative capacitance induced by a train of 10 depolarizations from -70 mV to 0 mV (gray trace) from control and *PIP4P2* knockdown human β-cells. **C.** Averaged cell size (n = 24 and 20), early exocytosis (n =24 and 20), late exocytosis (n = 24 and 20), total exocytosis (n = 24 and 20) obtained from **B** (n = 5 donors). **D.** representative current traces induced by a depolymerization from -70 mV to -10 mV (gray trace). **E.** Averaged $Na^+$ currents (n =24 and 20), early $Ca^{2+}$ currents (n = 24 and 20) and late $Ca^{2+}$ currents (n = 24 and 20) obtained from **D** (n = 5 donors). Data are presented as Mean ± SD. Student's t test. *** P < 0.001; ns – not significant.

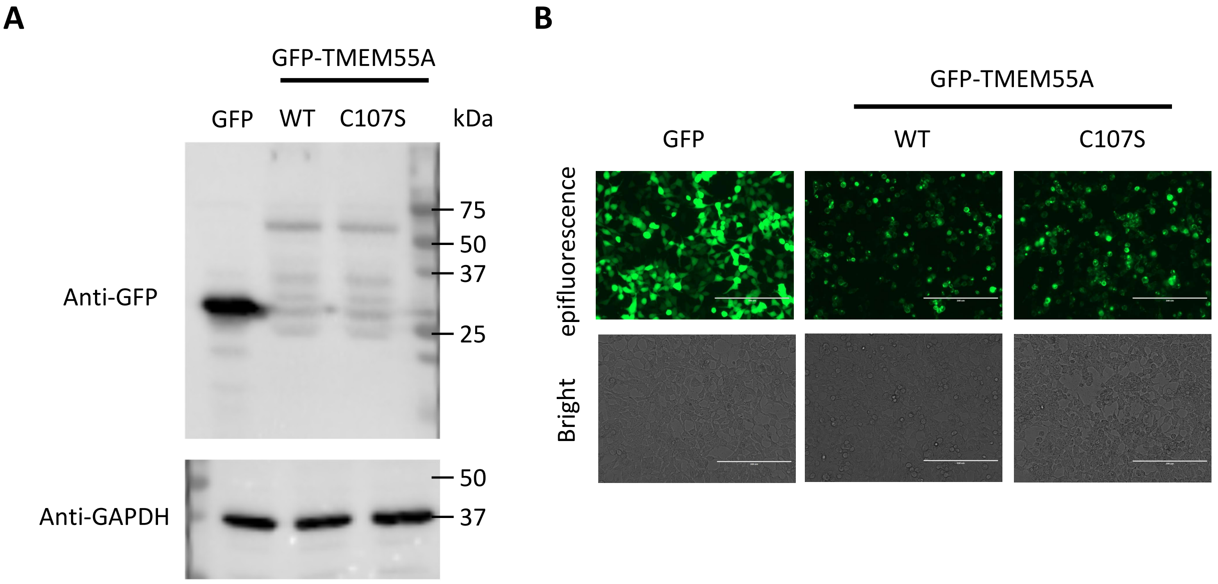


**Figure S2. Overexpression of GFP, GFP-TMEM55A, GFP-TMEM55A C107S in HEK cells.** **A.** Representative western-bolt shows the expression of GFP, GFP-TMEM55A, GFP-TMEM55A C107S in HEK cells. **B.** Epifluorescence and bright field images shows the expression of GFP, GFP-TMEM55A, GFP-TMEM55A C107S in HEK cells. Scale bar, 200 µm.

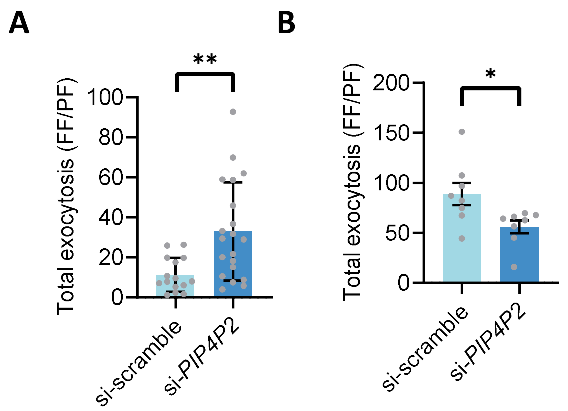


**Figure S3. Opposite effect of TMEM55A in α-cells from human T2D and mouse with HFD. A.** Averaged total exocytosis (n = 15 and 20 from 3 donors with T2D). **B.** Averaged total exocytosis (n = 8 and 8) obtain from 2 mice fed with HFD). Data are presented as mean ± SD. Student's t test. * $P < 0.05$; ** $P < 0.01$.

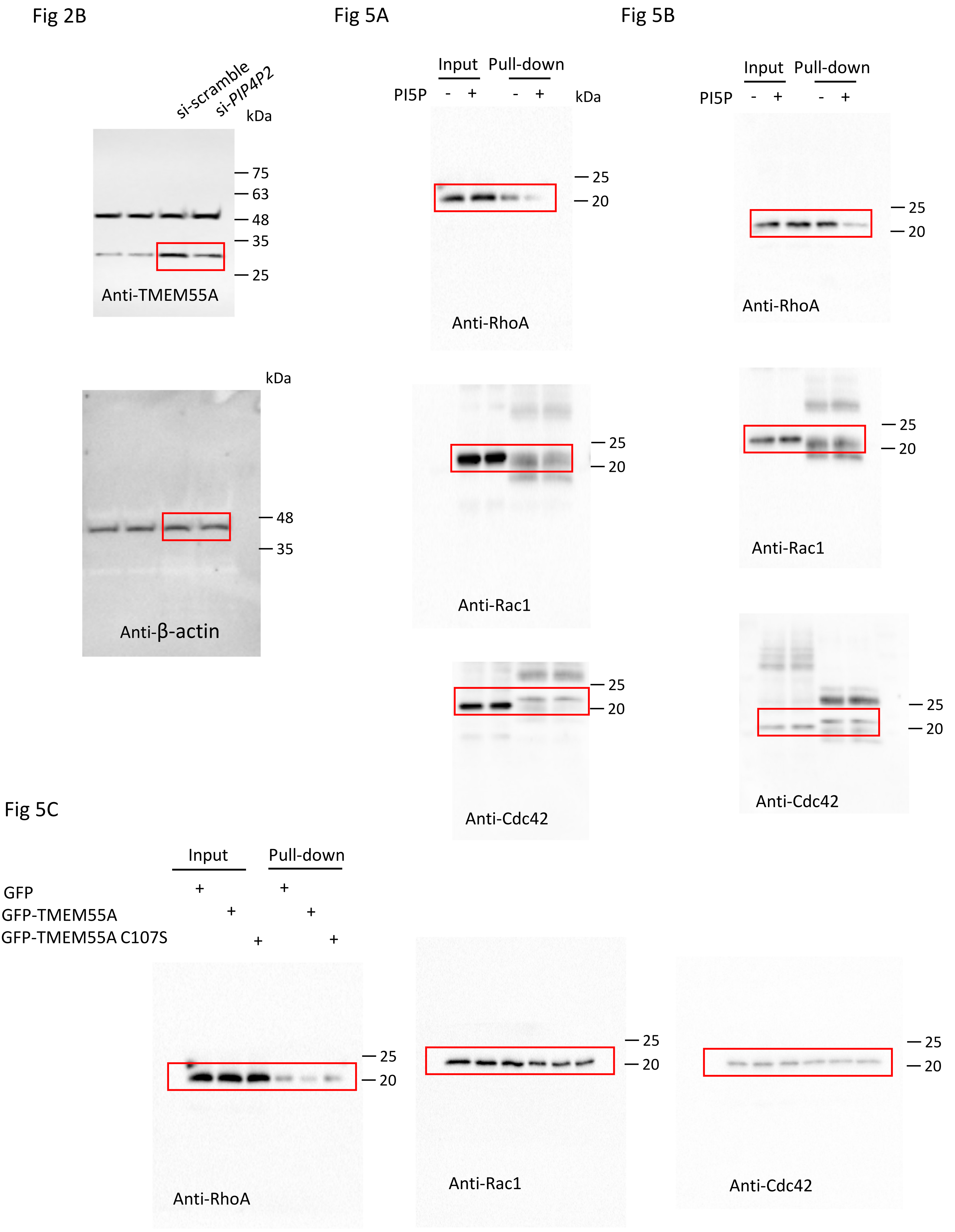


**Figure S4.** Uncropped scans of immunoblots included in main figures.

| | Forward primer (5’-3’) | Reverse primer (5’-3’) |
|---|---|---|
| **qPCR** | | |
| **Human PIP4P2** | GATGCCCAAGACCCAACTGTAGACG | CCACACACGACCCTTGTACCTTCTG |
| **Human PPIA** | TGACTTCACACGCCATAATGGCACTGG | AACACCACATGCTTGCCATCCAACCAC |
| **Mouse PIP4P2** | GATGTCCGAGACCCAACTGTCGACG | CCGCACACTACCCTTGTGCCTTC |
| **Mouse PPIA** | TGGCTATAAGGGTTCCTCCTTTCACAG | GCCAGGACCTGTATGCTTTAGGATG |
| **Subclone** | | |
| **Xhol-GFP-TM55AF** | ccactcgagATGGCTGCTGATGGG | - |
| **BamHI TM55AR** | - | ccaggatccTTATGCAAAACTGTGTTCTGGATAA |
| **Site-directed mutation and sequencing** | | |
| **TMEM55A C107S** | TCTTCTCATTAGTAAGGACACAT | CAATTACAAGGGCATCTAAC |
| **T7 promoter** | TAATACGACTCACTATAGG | - |

**Table S1.** Primers used in the current study for qPCR, subclone, site-directed mutation and sequencing.

| Target | Supplier | Catalog Number | Working dilution |
|---|---|---|---|
| **Primary antibodies** | | | |
| insulin | Dako/Agilent | IR002 (IR00261-2) | 1:5 for IF |
| glucagon | Sigma | G2654 | 1:200 for IF |
| TMEM55A | St John's laboratory | STJ195771-100 | 1:1000 for WB; 1:200 for IF |
| β-actin | Santa Cruz Biotechnology | sc-81178 | 1:1000 for WB |
| GAPDH | Santa Cruz Biotechnology | sc-137179 | 1:1000 for WB |
| Rac1 | Cytoskeleton, Inc | ARC03 | 1:1000 for WB |
| Cdc42 | Cytoskeleton, Inc | ACD03 | 1:1000 for WB |
| RhoA | Cytoskeleton, Inc | ARH03 | 1:1000 for WB |
| GFP | abcam | ab6556 | 1:1000 for WB |
| **Secondary antibodies** | | | |
| ECL Anti-Rabbit IgG, HRP | GE Healthcare | NA934V | 1:5000 for WB |
| ECL Anti-Mouse IgG, HRP | GE Healthcare | NA931V | 1:5000 for WB |
| Alexa Fluor 488 goat anti-gunia pig IgG (H+L) | ThermoFisher | A11073 | 1:200 for IF |
| Alexa Fluor 568 Goat anti-mouse IgG (H+L) | ThermoFisher | A11004 | 1:200 for IF |
| Alexa Fluor 647Goat anti-Rabbit IgG (H+L) | ThermoFisher | A-21244 | 1:200 for IF |

**Table S2.** Antibodies used in the current study.
